# Can root-associated fungi mediate the impact of abiotic conditions on the growth of a High Arctic herb?

**DOI:** 10.1101/2020.06.20.157099

**Authors:** Magdalena Wutkowska, Dorothee Ehrich, Sunil Mundra, Anna Vader, Pernille B. Eidesen

## Abstract

Arctic plants are affected by many stressors. Root-associated fungi are thought to influence plant performance in stressful environmental conditions. However, the relationships are not transparent; do the number of fungal partners, their ecological functions and community composition mediate the impact of environmental conditions and/or influence host plant performance? To address these questions, we used a common arctic plant as a model system: *Bistorta vivipara.* Whole plants (including root system) were collected from nine locations in Spitsbergen (n=214). Morphometric features were measured as a proxy for performance and combined with metabarcoding datasets of their root-associated fungi (amplicon sequence variants, ASVs), edaphic and meteorological variables. Seven biological hypotheses regarding fungal influence on plant measures were tested using structural equation modelling. The best-fitting model revealed that local temperature affected plants both directly (negatively aboveground and positively below-ground) and indirectly - mediated by fungal richness and the ratio of symbio- and saprotrophic ASVs. Fungal community composition did not impact plant measurements and plant reproductive investment did not depend on any fungal parameters. The lack of impact of fungal community composition on plant performance suggests that the functional importance of fungi is more important than their identity. The influence of temperature on host plants is therefore complex and should be examined further.

## Introduction

Arctic plants are facing many environmental constraints for growth, such as short vegetation season, consistent cold, limitation of nutrients or cyclic physical disturbances, i.e. cryoturbation^1^. These plants have evolved a range of adaptations to cope with the prevailing conditions, including being perennial and allocating most of their biomass below-ground^2–4^. Being perennial provides a resource-saving advantage in nutrient-poor habitats with low temperatures that slow down biochemical reactions and therefore also growth, whereas the benefits of biomass allocation to below ground parts include increased area of nutrient absorption. Because of nutrient scarcity, the interface between plant and soil is of relatively greater importance in the Arctic than in other biomes^3^. A significant part of the soil-plant interface is inhabited by microbes, including roots-associated fungi (RAF). Arctic RAF consist mostly of symbiotrophic fungi, especially ectomycorrhizal fungi^5–7^. These fungi efficiently increase the volume of soil that can be penetrated in search for resources, such as nutrients from seasonally or newly thawed permafrost^8^. The most severe limitations for growth observed in arctic plants are due to low temperatures and resource limitation^1,9^, suggesting that the relationship with RAF might play a crucial role in plant survival and growth.

Multiple characteristics of species communities play an essential role in the functioning of ecosystems, such as richness, abundance or community structure^10–13^. Based on previous findings, we may expect that the more diverse the community of RAF, the better for a host plant^14^. However, it is not clear how these characteristics of RAF communities impact their host plants, especially in cold biomes. Symbiotic fungi provide resources and probably additional benefits mitigating possibly harmful effects of environmental stressors enhancing plant growth and productivity^15^. However, releasing root exudates of primary metabolites that can be absorbed by members of its microbiome does come with a cost for a plant^16,17^. In nitrogen-limited tundra in Alaska, 61-88% of plant nitrogen (N) was supplied from mycorrhizal fungi. In exchange, the plants delivered 8-17% of carbon (C) produced photosynthetically to the fungi^18^. A plant could perhaps increase the amount of released nutritious root exudates to attract more species of symbiotrophic fungi that in turn, could potentially increase the amount of nitrogen delivered. However, higher fungal richness would increase competition for limited space in the rhizosphere and possibly for resources, although the mechanism is not yet fully described. Therefore, plants ‘living on the edge’ in the High Arctic may benefit from the selective choice of their members of RAF communities, favouring the most beneficial fungal partners for plant growth or mediation of stressors^19^. In this scenario, species richness in RAF communities would be irrelevant for plant performance. The presence of specific functional traits rather than their identity could be more important^20^. The vast array of interconnected biotic and abiotic factors occurring in natural systems complicate uncovering if and how plants show preference among their root-associated fungi among the pool of species present in the soil^21^.

One approach to disentangle these often confounded factors are controlled experiments. Most of the experiments assessing the impact of RAF diversity on host plant performance have focused on arbuscular mycorrhiza in crops^22^; whereas similar studies on ectomycorrhizal (EcM) plant species come mostly from the pre-high throughput sequencing era and have focussed on trees (e.g. ^14^). Several experiments under controlled settings have shown that EcM host plants may clearly benefit from their increased fungal richness, however, the tested level of richness was often incomparable with natural environments, such as an increase from 1 to 4 species of EcM fungi^14^. Some studies, however, did not find any enhancements in plant performance mediated by EcM fungi or concluded that the outcome of EcM species richness on plant productivity is context dependent^23^. RAF diversity was shown to be particularly sensitive to experimental conditions compared to fungi that inhabit space further from the roots in the rhizosphere or bulk soil^24^. Moreover, morphology and physiology of lab-grown plants differ from those in the natural systems, e.g. by increasing growth rate and higher concentrations of nutrients in tissues^25^. All these differences could affect and alter plant-associated organisms, such as RAF. Experimental procedures cannot consider all the complexity of natural systems and their effects do not always reflect those observed in the wild. Thus, observational studies can provide crucial complementary knowledge, in particular for extreme environments like the high Arctic.

Species response to environmental shifts, including ongoing climate changes, is one of the crucial questions in natural sciences. It is a particularly outstanding issue in the Arctic where rates of temperature and precipitation are changing at the fastest pace in the world, and are predicted to continue rising rapidly^26,27^. These changes impact mechanisms that alter biogeochemical cycles and determine critical ecosystem-climate feedback processes, such as the release of organic carbon of which nearly half of the global stock is stored in the Arctic soils^28,29^ or increased growth of vascular plants. Such ecosystem feedbacks, which are essential bricks in the understanding of global change, depend on complex relationships between abiotic and biotic factors in arctic soils^30^. However, the biology of these soils remains at present an understudied ‘black box’.

To shed some light onto these soil processes, we used a plant-centric approach to study the impact of the root-associated fungal community on the growth and reproductive investment of a wide-spread arctic plant, *Bistorta vivipara*. We took into account the most important abiotic factors which likely affect the host plant and its RAF community. We used structural equation modelling (SEM) to assess whether the fungal community mediates the effect of abiotic conditions on plant performance and to disentangle direct from indirect effects. We tested the following hypotheses: (i) Plant morphological measurements (considered as a proxy for plant performance) depend both on abiotic conditions and on the fungi community, and (ii) only richness and functional traits, but not the specific species composition of the RAF community affects plant morphology. Moreover, we tested, which measurements of plant parts involved in different processes such as energy storage, energy acquisition and reproduction depended on the RAF community.

## Methods

### Study system

To test our hypotheses, we selected alpine bistort *Bistorta vivipara* (L.) Delarbre (*Polygonaceae*), a model plant to study root-associated microbial communities in alpine^6,31–35^ and arctic habitats^7,19,36–40^. *Bistorta vivipara* is a common, long-lived perennial herb in the northern hemisphere. Its compact root system, combined with the ability to inhabit a range of habitats, makes this species a perfect candidate to study root-associated communities concerning environmental gradients, such as chronosequences^6,38,39^ or climate gradients^37^.

### Datasets

We combined and reanalysed datasets spanning over nine different locations in Spitsbergen, the largest island of the high-arctic archipelago Svalbard, Norway (Table 1; Figure 1). Each dataset consisted of host morphology, molecular descriptions of the RAF community, together with associated edaphic variables (Table 1). Each of the studies established a randomized sampling scheme in the locality of choice, also assuring that sampled plants are of different age. Whole plants with an intact root system were excavated. To explore the associations between plant performance, allocation patterns and its environment we measured three morphological features of the *B. vivipara* individuals hosting the analysed RAF communities (Supplementary 1): The rhizome is an underground storage organ that accumulates assimilated biomass as nonstructural carbohydrates, therefore here we used it as a proxy for overall plant performance^41^. Rhizome dimensions were measured and used to calculate an approximate volume (RV) by multiplying its length, height and width. Length of the longest stem leaf (LL) was used as a proxy for photosynthetic capabilities of the plant – the longer the leaf, the bigger photosynthetic area. In the upper part of the stem, *B. vivipara* produces flowers and bulbils for sexual and asexual reproduction, respectively. We used the ratio of the length of the stem covered by flowers and bulbils (inflorescence) to the total stem length (I/S), as a proxy for the plant’s investment in reproduction.

**Figure 1.**
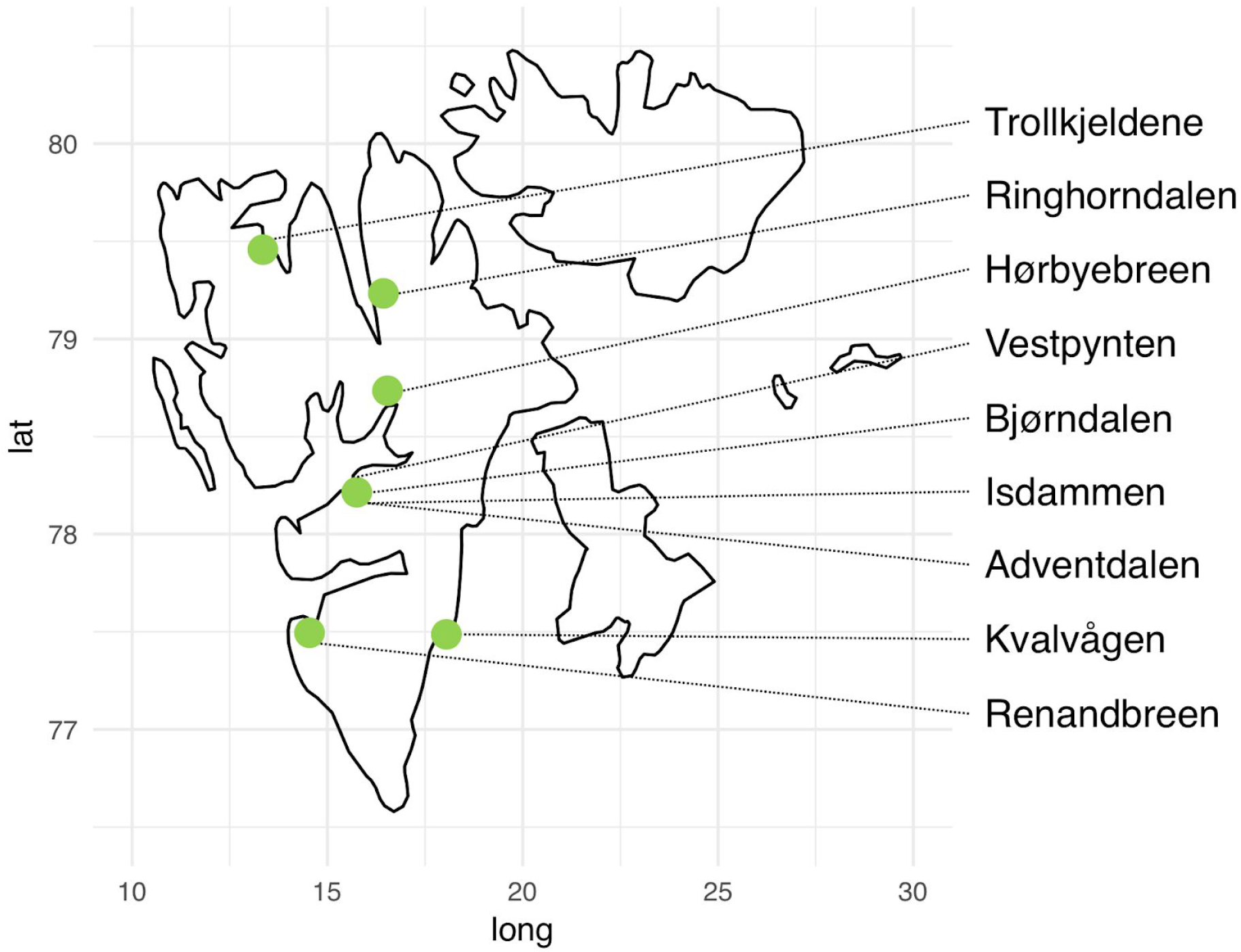
*Bistorta vivipara* plants from the four concatenated datasets were collected in nine localities on Spitsbergen.

**Table 1.**
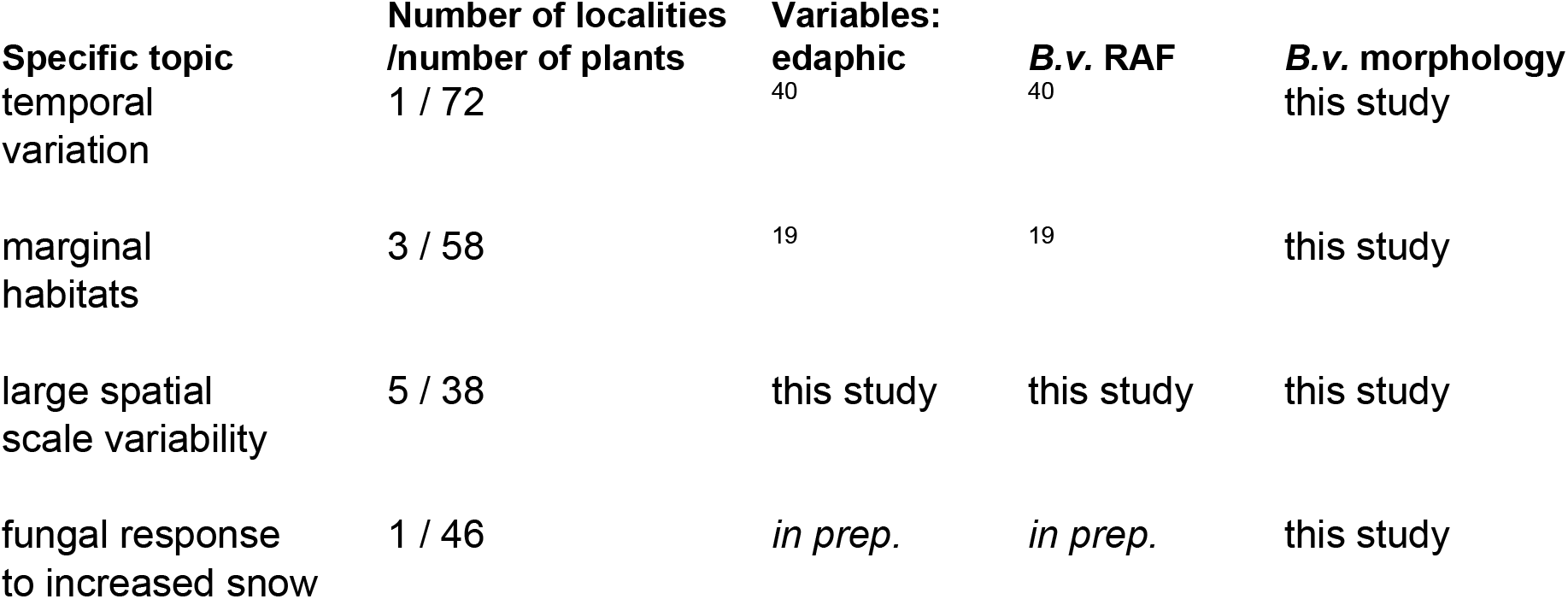
Overview of the data included in this study. Each dataset was generated to investigate specific topics regarding *Bistorta vivipara* root-associated fungi (RAF). References are given for previously published data.

### Meteorological and edaphic variables

Meteorological data were obtained for each sampling point from the high-resolution 1 km-gridded dataset Sval_Imp_v1^42^. We extracted the sum of average monthly precipitation (p) and average July air temperature (t), both from the year of sampling.

Soil samples were collected from the same sampling spot as plants. The following edaphic parameters, representing critical properties of the abiotic environment, were measured in all datasets: pH, soil nitrogen concentration (N) and carbon to nitrogen ratio (C/N; used as an indicator for soil nitrogen availability or soil fertility). Edaphic variables were obtained in the same way for all datasets (described in detail in^7,19,40^).

### Fungal data

*Bistorta vivipara* roots were cleaned within a day from sampling and fixed in a 2% CTAB extraction buffer until DNA extraction (details described in each of the publications; Table 1). All datasets targeted the same fragment of internal transcribed spacer 2 amplified with fITS7a forward primer^43^ and reverse primer ITS4^44^ and sequenced with Illumina MiSeq (300bp paired-end reads).

Each dataset was a mixture of sequences located in ‘forward’ and ‘reverse’ direction. Thus, first, a mapping file with variable length barcodes and primer sequences was used to identify sequences in each location using sabre (https://github.com/najoshi/sabre) and generating separate R1 and R2 files for each read direction. Next, primers were clipped, and sequences with ambiguous bases (Ns) were removed using cutadapt v. 2.5^45^. Python script FastqCombinePairedEnd.py (https://github.com/enormandeau/Scripts) was used to assure that each sequence had its pair and were in the matching order for further analyses. We used an amplicon sequence variants (ASVs) approach implemented in DADA2 v. 1.11.1^46^ and executed in R v. 3.5.2^47^ (for details see Supplement 2 and scripts generated for this study). The datasets were analysed using DADA2 ITS workflow (https://benjjneb.github.io/dada2/ITS_workflow.html). Fungal data were produced independently for each study; therefore, they were initially analysed separately due to different error rates for each sequencing run. Separate ASVs tables were then merged. Consensus method was used to remove chimaeras (3759 out of 11243 input sequences). Sequences shorter than 200bp and six samples with a very low number of reads were removed. Due to profound differences in depth of sequencing the ASV table was randomly subsampled (21639 reads per sample; number of detected ASVs before and after subsampling was highly correlated; Kendall’s ⊺ = 0.95). Taxonomy was assigned using the RDP naive Bayesian classifier implemented in DADA2 with a full UNITE+INSD reference dataset for fungi^48^ (sh_general_release_dynamic_02.02.2019). All the ASVs were functionally annotated using the FUNGuild database^49^.

Differences in community composition were summarized through non-metric multidimensional scaling (GNMDS; *vegan* package^50^), and we used the first axis as a proxy for composition in further analyses. We used both presence-absence based metrics and parameters based on read abundance to describe RAF communities: ASV diversity (D), a ratio of symbio- to saprotrophs (Sy/Sa) and GNMDS values for 1^st^ axis as a proxy for community composition (CC; Table 2).

**Table 2.** Metrics used to describe the fungal community used in this study for presence absence data and number of reads, respectively. All the parameters were calculated using a rarefied table containing amplicon sequence variants (ASVs).

| Fungal parameter | Presence-absence table | Abundance table |
| --- | --- | --- |
| Diversity (D) | richness<br>(number of ASV) | Shannon-Wiener (H') index |
| $\frac{\text{Symbio-}}{\text{Saprotrophs}}$ (Sy/Sa) | ratio of ASVs | ratio of reads |
| Community<br>composition (CC) | GNMDS 1<sup>st</sup> axis score | GNMDS 1<sup>st</sup> axis score |

### Statistical analyses and model selection

The statistical analyses were performed in R v. 3.5.2^47^. Based on available literature of soil and weather influence on fungi and plant interactions in the Arctic (Table 3), we built seven hypothetical causal path models relating abiotic variables to the three metrics characterizing the fungal community and plant morphological measurements (solid lines in Figure 2). The unbranched rhizome of *B. vivipara* elongates with age, providing space for new roots to stem from its distal end^51^ and therefore increasing the richness of recruited RAF^34^. Randomised sampling schemes in each of the studies included in our study excluded the potential influence of plant age on the results. For the full model, we assumed that all three fungal parameters influence all three plant measures, additionally to abiotic factors impacting both fungal and plant variables.

**Figure 2.**
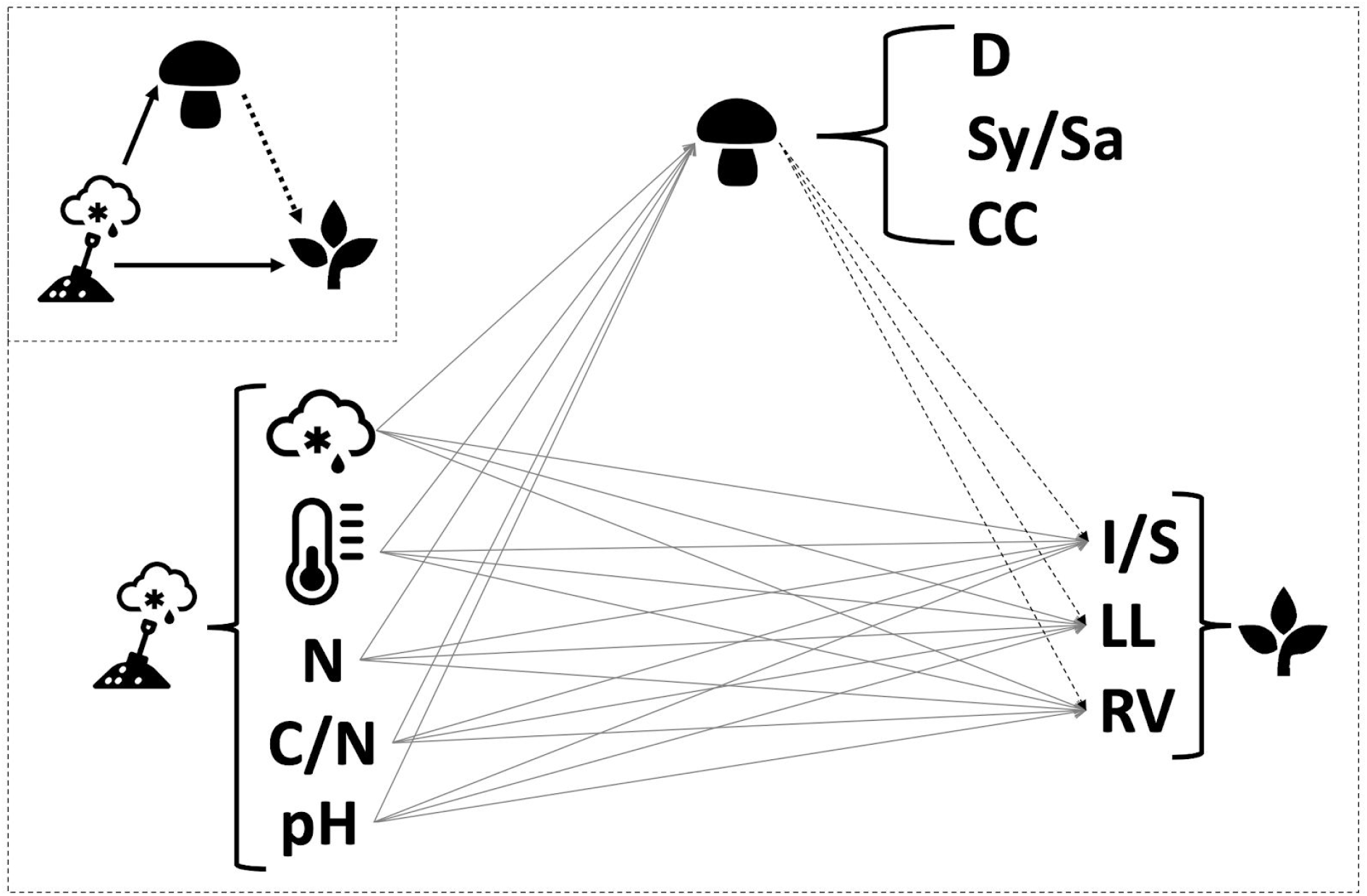
Schematic illustration of a conceptual plant-centric model representing relationships between variables suggested by the literature and tested in this study. Solid lines are associations were researched by studies from the Arctic; dashed lines were described by fewer studies, mainly from other regions. The full model includes all possible links between each abiotic, fungal and plant variable. Abbreviations and symbols: N - soil nitrogen content; C/N - the ratio of soil nitrogen to soil carbon content; p- precipitation; t - temperature; D - diversity ; Sy/Sa - the ratio of symbio- to saprotrophs; CC - fungal community composition; I/S - the ratio of inflorescence to stem length; RV - rhizome volume; LL - leaf length of the longest leaf.

**Table 3.**
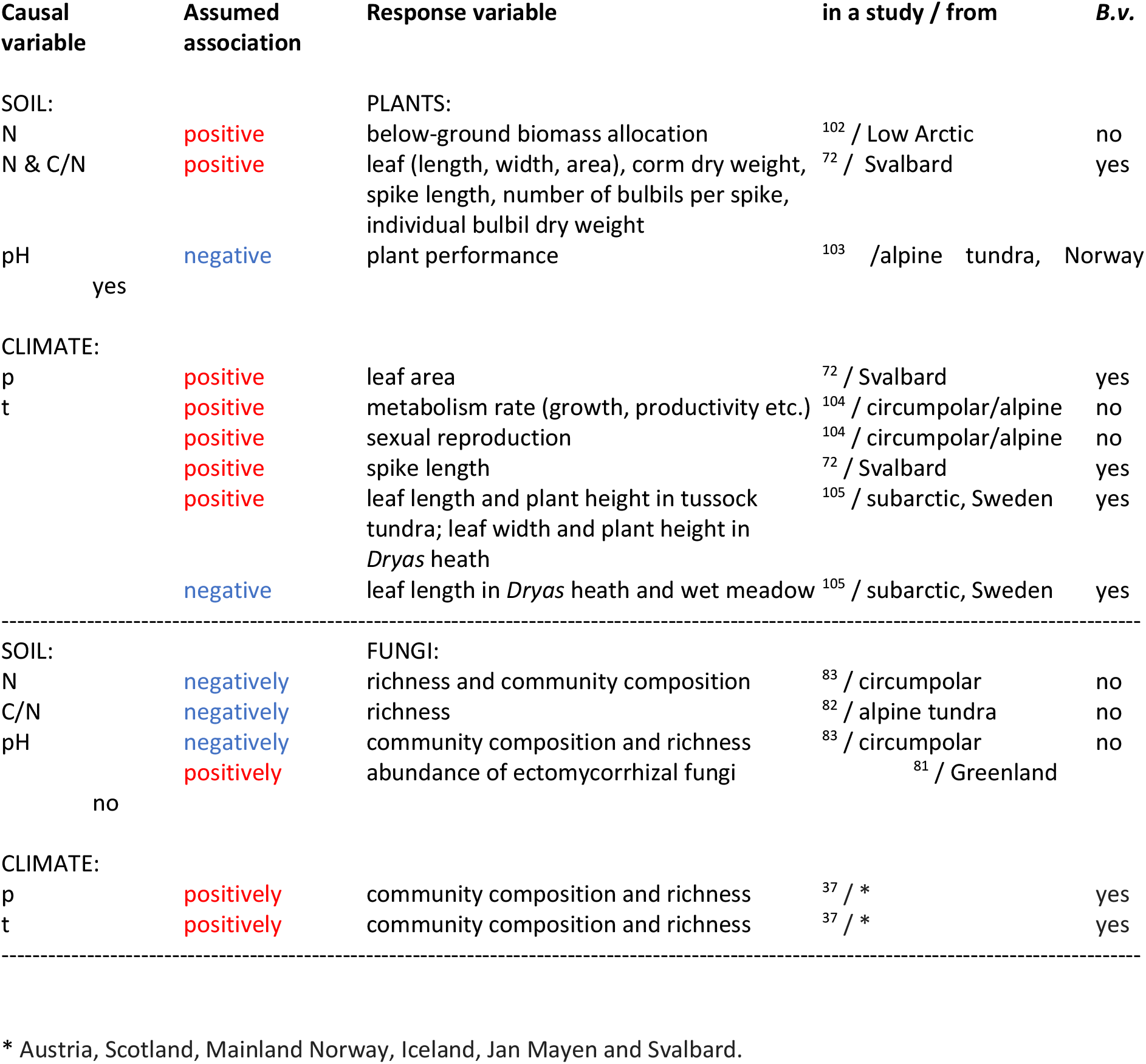
Relationships between abiotic factors and root-associated fungi or plant metrics documented in the literature. Some of the relationships have been demonstrated generally for arctic plants and arctic fungi, and have not been specifically shown in *B. vivipara*. Abbreviations: N - soil nitrogen content; C/N - ratio of soil nitrogen to soil carbon content; p - precipitation, t - temperature, *B.v*. - whether the study was specifically conducted on *B. vivipara* plants or *B. vivipara* root-associated fungal communities.

| Causal variable | Assumed association | Response variable | in a study / from | *B.v.* |
| --- | --- | --- | --- | --- |
| SOIL: |  | PLANTS: |  |  |
| N | positive | below-ground biomass allocation | <sup>102</sup> / Low Arctic | no |
| N & C/N | positive | leaf (length, width, area), corm dry weight, spike length, number of bulbils per spike, individual bulbil dry weight | <sup>72</sup> / Svalbard | yes |
| pH | negative | plant performance | <sup>103</sup> / alpine tundra, Norway |  |
| yes |  |  |  |  |
| CLIMATE: |  |  |  |  |
| p | positive | leaf area | <sup>72</sup> / Svalbard | yes |
| t | positive | metabolism rate (growth, productivity etc.) | <sup>104</sup> / circumpolar/alpine | no |
|  | positive | sexual reproduction | <sup>104</sup> / circumpolar/alpine | no |
|  | positive | spike length | <sup>72</sup> / Svalbard | yes |
|  | positive | leaf length and plant height in tussock tundra; leaf width and plant height in *Dryas* heath | <sup>105</sup> / subarctic, Sweden | yes |
|  | negative | leaf length in *Dryas* heath and wet meadow | <sup>105</sup> / subarctic, Sweden | yes |
| SOIL: |  | FUNGI: |  |  |
| N | negatively | richness and community composition | <sup>83</sup> / circumpolar | no |
| C/N | negatively | richness | <sup>82</sup> / alpine tundra | no |
| pH | negatively | community composition and richness | <sup>83</sup> / circumpolar | no |
|  | positively | abundance of ectomycorrhizal fungi | <sup>81</sup> / Greenland |  |
| no |  |  |  |  |
| CLIMATE: |  |  |  |  |
| p | positively | community composition and richness | <sup>37</sup> / * | yes |
| t | positively | community composition and richness | <sup>37</sup> / * | yes |

\* Austria, Scotland, Mainland Norway, Iceland, Jan Mayen and Svalbard.

Next, we hypothesized that fungi might not be essential for specific plant measurements. Therefore, in the three subsequent models, we preserved all the relationships omitting only the fungal variables in a specific plant response (I/S, RV or LL does not depend on fungi). In the next models, we, therefore, hypothesized that CC is not an important parameter for any of the plant measurements. Additionally, we combined this last model with the best model obtained from simplifying the relationships between fungi and plants responses.

Finally, to evaluate whether fungal parameters have any impact on plant measurements, we removed all connections between fungal parameters and plant measurements. In the models, we treated edaphic and meteorological variables as independent. We are aware that they can affect each other, but this was not the focus of the study. The most considerable correlation among them was between N and C/N (r = −0.64). We did also not hypothesize any causal links between the fungal parameters. Concerning the plant variables, we assumed a causal link between rhizome volume and leaf length, because leaf growth in the start of the season depends on stored resources. Locality was used as a random effect in all the models because fungal community composition usually shows a high spatial variation (e.g.^36^) and because preliminary ordinations showed that in our dataset fungal communities differed between localities.

We applied structural equation modelling (SEM) to carry out an exploratory path analysis of these models, using the psem function in the piecewiseSEM package^52^. The SEM was composed of linear mixed-effects models (LMMs) for each fungi parameter and plant measurement, which was fitted using the lme function in *nlme* package^53^. The fit of the separate LMMs were assessed graphically for normality of the residuals. Residuals clearly deviating from the expected distribution on a quantile-quantile plot with standardised residuals > |3| were considered as outliers and therefore excluded.

The analysis was performed using both presence-absence based and read abundance metrics for the fungal community. Because some of the fungal parameters were correlated, we included non-directed correlations among them in the SEM to make it possible to estimate the paths in our exploratory model. It was the case for CC and Sy/Sa based on presence-absence and for Sy/Sa and D based on read abundance. The distributions of all variables were assessed graphically, and some were log- or logit-transformed to assure roughly normal distributions. All variables were scaled to 0 mean and a standard deviation of 1 to make effect sizes comparable.

A prerequisite for a SEM model to be considered as fitting was Fisher’s C p-value > 0.05^54^ The best models among the candidate sets described above were chosen based on the lowest AIC values. Both of these values were calculated within the *psem* function. We used statistically significant estimates from the best fitting presence-absence model to calculate indirect effects of abiotic factors on plant measures.

The combined dataset consisted of 214 *B. vivipara* plant measurements with associated edaphic data and corresponding RAF data. For the SEM, we excluded all observations with missing values resulting in a final dataset with 188 plants (after excluding outliers presence-absence dataset had 187 and abundance dataset 185 values).

## Results

### Models based on presence-absence fungal parameters

The best-fitting presence-absence path model (AIC_min_ = 117.97; Table 4) supported the hypothesis that fungal CC does not impact plant measurements, and simultaneously no fungal parameters affect the I/S. The second best-fitting model with a relative difference ΔAIC < 1, supported a related hypothesis that I/S does not depend on any fungal parameters included in this study, but included the effect of CC on other plant parameters.

**Table 4.** Summary of the models and statistics used for best fitting model selection. Each model reflects a separate hypothesis. The full model includes all possible links between each fungal variable and each plant variable. Subsequent models exclude some of the links, as indicated in the name of each model. Abbreviations: I/S - ratio of inflorescence to stem length; RV - rhizome volume; LL - leaf length; CC - root-associated fungal community composition.

| **Model** | **Fisher's C** | **p** | **AIC** |
| --- | --- | --- | --- |
| <u>Presence-absence</u> |  |  |  |
| Full | 3.2 | 0.780 | 121.23 |
| I/S does not depend on fungi | 8.9 | 0.837 | 118.86 |
| *RV does not depend on fungi* | *28.0* | *0.014* | *138.00* |
| LL does not depend on fungi | 22.3 | 0.073 | 132.31 |
| Fungal CC not important | 10.3 | 0.739 | 120.32 |
| **Fungal CC not important + no I/S** | **12.0** | **0.849** | **117.97** |
| *No effect of fungi on plants* | *42.9* | *0.020* | *140.86* |
| <u>Abundance</u> |  |  |  |
| Full | 7.1 | 0.529 | 123.07 |
| I/S does not depend on fungi | 11.7 | 0.632 | 121.68 |
| RV does not depend on fungi | 13.6 | 0.482 | 123.58 |
| LL does not depend on fungi | 13.4 | 0.492 | 123.44 |
| Fungal CC not important | 13.5 | 0.491 | 123.45 |
| Fungal CC not important + no I/S | 13.5 | 0.759 | 119.53 |
| **No effect of fungi on plants** | **21.3** | **0.728** | **119.28** |

The best model for each approach is highlighted in **bold**.
Models which don't fit based on the test of directed separation are in *italics*.

In the best-fitting and most parsimonious model, fungal community richness and the ratio of symbiotrophic to saprotrophic species were related to plant measurements as follows (Figure 3a): fungal richness with RV (positive path coefficient (PC ± SE = 0.26 ± 0.07, p < 0.001); full list of all the effect sizes in Supplement 4a) and Sy/Sa with LL (PC ± SE = −0.20 ± 0.07, p = 0.004). Except for the fungal metrics, the RV also showed positive correlations with p (PC ± SE = 0.29 ± 0.11, p = 0.01). LL was negatively impacted by N content (PC ± SE = −0.20 ± 0.08, p = 0.02) and t (PC ± SE = −0.34 ± 0.08, p < 0.001). The highest estimate in our model suggested correlation between RV and LL (PC ± SE = 0.53 ± 0.06, p < 0.001).

**Figure 3.**
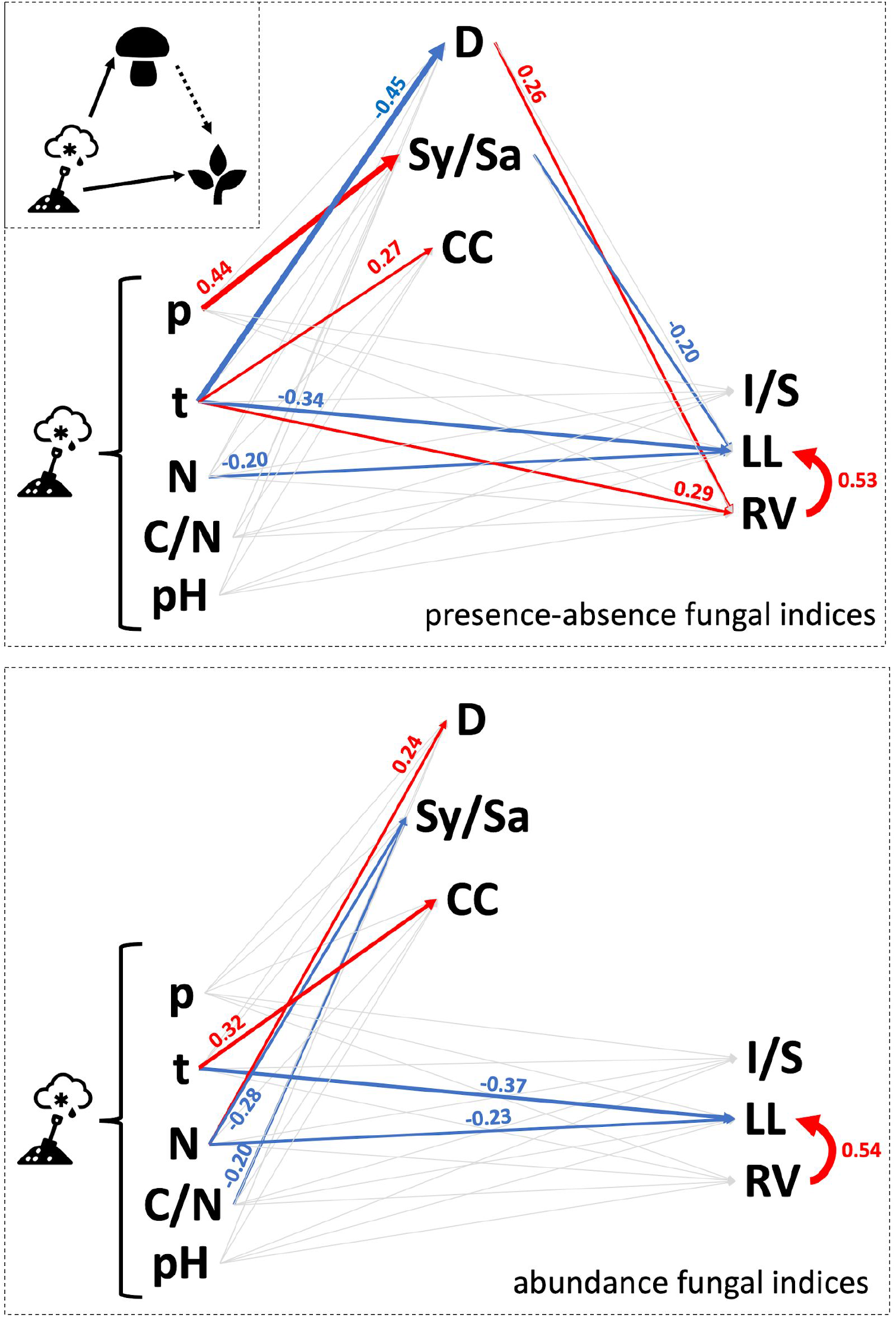
Path diagram showing tested connections between predictor and response variables in the best fitting models. Statistically significant (p < 0.05) links are depicted by arrow colours (positive or negative nature of the relationship) and thickness (relationship magnitude); the numbers are estimates from the models. Abbreviations and symbols: N - soil nitrogen content; C/N - the ratio of soil nitrogen to soil carbon content; p - precipitation; t - temperature; D - diversity ; Sy/Sa - the ratio of symbio- to saprotrophs; CC - fungal community composition; I/S - the ratio of inflorescence to stem length; RV - rhizome volume; LL - leaf length of the longest leaf.

Meteorological data had a clear effect on fungal parameters: p with Sy/Sa (PC ± SE = 0.44 ± 0.21, p < 0.04), and t with fungal CC (PC ± SE = 0.27 ± 0.09, p = 0.003) and D (PC ± SE = −0.45 ± 0.13, p < 0.001). Based on the best fitting presence-absence model, edaphic variables did not seem to impact any of fungal parameters and plant measurements except the already mentioned N content impact on LL. On the other hand, t correlated with multiple fungal and plant variables.

Among abiotic factors impacting plant measurements, t affected LL over three pathways: direct (negative, PC = −0.34) and two indirect: positive through RV (PC = 0.29 * 0.53 = 0.154) and negative through fungal D (PC = −0.45 * 0.26 = −0.117). The direct effect was therefore the strongest and the two indirect effects were of comparable magnitude, but opposite directions.

### Abundance model

The best-fitting path model based on read abundance supported the hypothesis that fungal parameters do not impact any plant measurements (AIC_min_ = 119.28; Table 4). Another model that differed by ΔAIC = 0.25 supported the same hypothesis as the best fitting presence-absence model: fungal CC does not impact plant measurements, and I/S is not affected by other fungal parameters either.

Although the role of fungi in the best model differs fundamentally from the best model based on presence-absence ASV table, they both preserved some of the same statistically significant relationships between environmental variables and plant measurements (Figure 3b, full list of all the effect sizes in from both types of models in Supplement 4). This included correlations between N content and LL (PC ± SE = −0.23 ± 0.08, p = 0.005), t and LL (PC ± SE = −0.37 ± 0.08, p < 0.001), as well as t and CC (PC ± SE = 0.31 ± 0.09, p < 0.001). Also, the relationship between two plant variables, RV and LL, showed the same magnitude as in the best fitting presence-absence model (PC ± SE = 0.54 ± 0.06, p < 0.001). This model supported no indirect effects of abiotic factors mediated by fungal parameters.

The abundance-based model revealed links between edaphic and fungal parameters that were not statistically significant in the presence-absence model. N content and C/N ratio correlated negatively with Sy/Sa (PC ± SE = −0.28 ± 0.10, p = 0.007 and PC ± SE = −0.20 ± 0.10, p < 0.04; respectively). The N content positively impacted fungal diversity (PC ± SE = 0.24 ± 0.11, p < 0.04).

### Variance in fungal and plant response variables

In both best fitting models, the variance in plant measurements was on average better explained by fixed factors than the variance in fungal parameters (marginal R^2^ = 0.02-0.44 vs 0.07-0.26, Table 5). However, overall the variance explained by fixed factors was rather low. On the contrary, locality included as a random factor explained on average more variation in fungi than in plants (conditional R^2^ - marginal R^2^ = 0.03 - 0.58 and 0.01 - 0.33, respectively). The high proportion of variance explained for fungal response variables was especially pronounced in presence-absence compared to the abundance model (conditional R^2^ - marginal R^2^ = 0.40 - 0.58 and 0.03-0.48, respectively).

**Table 5.** Proportion of variance explained without (marginal R^2^) and with random factors (conditional R^2^). Locality was used as a random factor in all of the models. Abbreviations: D - diversity; Sy/Sa - the ratio of symbio- to saprotrophs; CC - fungal community composition; I/S - the ratio of inflorescence to stem length; RV - rhizome volume; LL - leaf length of the longest stem leaf.

| Response | Presence-absence model<br>**CC does not impact plants + no I/S** |  | Abundance model<br>**No effect of fungi on plants** |  |
| --- | --- | --- | --- | --- |
| | Marginal $R^2$ | Conditional $R^2$ | Marginal $R^2$ | Conditional $R^2$ |
| Fungal: |  |  |  |  |
| D | 0.11 | 0.51 | 0.07 | 0.32 |
| Sy/Sa | 0.16 | 0.56 | 0.08 | 0.11 |
| CC | 0.26 | 0.84 | 0.18 | 0.66 |
| Plant: |  |  |  |  |
| I/S | 0.02 | 0.34 | 0.02 | 0.33 |
| RV | 0.24 | 0.39 | 0.15 | 0.24 |
| LL | 0.44 | 0.46 | 0.42 | 0.43 |

## Discussion

Establishing functional relationships between biological components, such as a host plant and its root-associated microbiome, taking into account abiotic drivers, could enhance the current understanding of soil carbon pools and decrease associated uncertainties^55,56^. To narrow these gaps, we studied the common arctic host plant *B. vivipara* and its RAF communities in connection with their environment. Here, we linked above- and below-ground plant measurements to fungal parameters, all assumed to be influenced by the same edaphic and meteorological conditions. This exploratory study revealed that measurements of below- and aboveground plant organs responded in opposite ways to temperature, the effects of which were both direct and mediated by parameters of the RAF community. Regarding fungal parameters, both species richness and functional diversity were important for plant performance measurements, but not the specific community composition.

Our study revealed that among the abiotic factors temperature was the most important for biotic elements, which reflects its immense significance in physical constraints for arctic biota^1^ and the general tendency of modifying interactions between organisms^57^. However, our results also suggest that the impact of temperature on an arctic host plant is far more complex than previously thought^58,59^ and in general, perhaps unpredictable^60^. The mechanism behind fungal mediation of temperature is not clear. Here we looked only into a few parameters associated with RAF communities that impacted the plant both positively and negatively balancing themselves out. However, there are other molecular and physiological characteristics that could explain the influence of fungi on plant performance mechanistically. For instance, secretion of fungal signalling molecules, such as volatile organic compounds^61^ or plant-like hormones^62,63^, that can be translocated to host plant cells and there elicit a physiological response. Release of these molecules could be temperature-dependent. Similarly, plant-based responses to these signals could also be at least partly temperature-dependent, e.g. release of root exudates^64^.

Different influences of temperature on below- and aboveground plant measurements could question current methods of monitoring changes in arctic vegetation, such as the normalized difference vegetation index (NDVI) used as a proxy for plant biomass. This technology advanced the understanding of vegetation biomass dynamics simultaneously over vast and otherwise under-sampled areas of the Arctic (e.g.^65–67^). However, it is based on remote measurements of Earth’s surface reflectance, and therefore takes into consideration only aboveground changes in foliage. In these methods plants’ below-ground productivity and biomass are omitted, probably resulting in underestimation of the overall impact of increased temperatures on plants, such as *B. vivipara*, which is an ubiquitous species in the Arctic and essential food source for ptarmigans^68^, geese^69^ and reindeer^70^. Temperature had a direct opposite effect of similar magnitude on LL and RV (−0.34 vs 0.29, respectively), additionally strengthened by indirect fungal effects, which suggests that NDVI can easily underestimate the impact of warming on overall plant biomass and misjudge understanding of carbon stocks dynamics. Presently, there are no tools that could be used to scan below-ground plant biomass at scales similar to NDVI. However, there are some more laborious *in situ* methods, e.g. minirhizotrons, that are used to measure below-ground biomass^71^. Their use significantly enhances our understanding of the dynamics in belowground biomass allocation. Nevertheless, the implications of temperature affecting a host plant through multiple pathways generate major difficulties in projections of the future response of ecosystems to warming.

Negative impact of nitrogen on leaf length was unexpected in the light of previous findings^72^. *Bistorta vivipara* is regarded as a pioneer plant^73^, able to cope with severe conditions and resource limitations^32,39^. In a High Arctic nitrogen-rich habitat, such as bird cliffs, where the competition between organisms is high, it is most likely outcompeted by other plants. Additionally, these highly nutritious habitats are characterised by an increased number of plant interactions with herbivores, such as reindeers, that can eliminate foliage.

Almost all symbiotrophic RAF of *B. vivipara* in Svalbard are ectomycorrhizal^7,39^. Since these fungi exchange nitrogen with plants in return for versatile carbon metabolites^18^, we hypothesized that in a resource-limiting environment this fungal trophic mode could promote bigger plants^74^, therefore bigger leaves. This way, fungi could potentially influence the number and amount of metabolites that the plant could produce in return and share in its rhizosphere. However, our results showed the opposite scenario, where Sy/Sa had a negative effect on leaf length, which suggests that more fungal partners enhance competition over resources that are scarce^75^. The richness of symbio- and saprotrophs taken into account separately did not show any associations with plant measurements (data not shown); however, the ratio of their richness did, perhaps reflecting the characteristics of soil conditions in different localities. Particularly small ratio of Sy/Sa was found in localities with little organic matter (Supplementary 3), suggesting that this parameter mirrors fertility properties of soil. When soil organic matter content is low, then colonizing plant roots ensures access to an easily accessible pool of carbon from root exudates^76^. Although *B. vivipara* root system is relatively compact and flexible, growing in mineral soils, including some stages of soil development of glacier forefronts^77^, could promote longer roots to assure access to quickly drained soil water. Intense disturbance caused by periglacial processes in these habitats may contribute to physical breaks in fine roots or associated fungal mycelium, perhaps leading to an increase in the number of saprotrophic species. Alternatively, saprotrophic fungi could be one of the first organisms in primary community assembly using organic carbon from previously unrecognized heterotrophic communities of invertebrates which feed on allochthonous organic matter now recognized as a crucial step of primary succession before establishment of autotrophs^78–80^.

Our finding that fungal community composition did not affect plant measurements could perhaps originate from strong environmental filtering on root-associated fungal communities^36^. High physicochemical heterogeneity of arctic soils corresponds with distinct RAF community composition observed at different scales^5–7,37^. On the one hand, a set of physicochemical conditions that translates into ecological niches selects species that can withstand and thrive in these locality-specific combinations of factors. Among them principally abiotic factors were shown to affect fungal parameters ^37,81–83^. Relationships between variables established based on the literature search (Table 3) were, in general, poorly reflected in the results of our models. In most cases, we saw no effects of abiotic drivers identified in the literature on neither plants nor fungi. It was especially pronounced in RAF community composition, suggesting other sources of the differences that are specifically connected to locality^19^. These could be other edaphic factors not included in this study (e.g. phosphorus^84^ or heavy metal concentrations^85^, competition^75,86^ or other factors that historically impacted the community assembly^87^. Nevertheless, the fact that arctic ectomycorrhizal RAF display little or no affinity to host species^88^ suggests that the fungal contribution to plants reflects mitigation of effects of locality-specific conditions, rather than individual species needs. Similar conclusions were made in edge soil habitats beyond the Arctic. For instance, RAF communities in soil characterised by combined effects of poor nutritional and water status^89^ or high contamination levels^90^ seem to also be host-independent and highly variable among the sites.

To explain discrepancies in results between presence-absence and read abundance models, it is necessary to identify possible sources of variation in read abundances in fungal metabarcoding studies. Fungal species vary in the copy number of ribosomal DNA (14-1442), and this number is independent of genome size or ecological roles, such as guild or trophic mode^91^. Strains of the same fungal species, especially yeast, can exhibit high variation of rDNA copy number^92,93^. Relative abundances of reads are sometimes used as a proxy for the relative biomass contributions of some species^94^. However, a quantitative meta-analysis found only a weak relationship between the two^95^. Read abundance can be profoundly affected by methodological biases at several steps during metabarcoding procedures, starting from the choice of primers through wet-lab methods, including sequencing, to bioinformatic pipelines ^96–99^. However, in our study, main pathways affecting plants directly and not through fungal parameters remained present in both best-fitting models. This supports prevalence of a biological signal over methodological biases from abundance data. On the other hand, the abundance-based model in this study showed clear links between fungal parameters and soil fertility (N and C/N) mirroring the stoichiometric state of the environment^100^ and temperature that controls the rate of biochemical reactions.

Here we demonstrated that fungal parameters, such as richness and functional diversity, could mediate the influence of abiotic factors on host plants, but it is not clear what are the mechanisms behind this. It is not clear how different fungi contribute to plants’ biometrics, how many resources are being exchanged with plants and how that changes with RAF variation in time and space. Not only molecular identification, but also establishing biomass estimations for both fungi and bacteria could help to understand below-ground dynamics. Low proportion of variance explained by fixed factors showed that there is a strong need to obtain and include more abiotic and biotic variables that were not considered in this study, but are of high importance for fungi and plants. Controlled experiments could potentially help to address these uncertainties. Additionally, morphological characterization of multiple plant species, biomass and nutrient concentration measurements in separate plant parts would ensure precise comparisons between plant life strategies in variable habitats and distant locations. Another critical aspect in making these links is to include the host plant genotype to tie its phenotype with the influence of the environment accurately^101^. A comprehensive interdisciplinary study employing various methods could help to develop a mechanistic understanding of links between above- and below-ground biota, including other taxonomic groups.

## Supporting information

Supplementary materials for Wutkowska et al., Can root-associated fungi mediate the impact of abiotic conditions on the growth of a High Arctic herb?

## Acknowledgments

The authors are thankful to all the people that contributed to the terrestrial part of MicroFun project, collected and processed the samples. This research was funded by University Centre in Svalbard, as well as ConocoPhillips and Lundin Petroleum through The Northern Area Program. The Svalbard Science Forum is acknowledged for providing Arctic Field Grant to SM (2012 and 2013), for sampling in Svalbard (project code 220126/E10; RIS ID 5009). We also thank the Governor (Sysselmannen, Longyearbyen) for permitting us to collect the root and soil samples from Svalbard.

## Author Contributions Statement

MW, DE, SM, PB - wrote and edited the manuscript; MW, AV - analysed sequencing data; MW, DE - did statistical modelling; SM - collected and processed soil samples, measured abiotic parameters, generated sequencing data, PB - developed the idea and

## Additional Information

The already published datasets are available online. All the other datasets used in this study are available at zenodo.com/addproperlink. Scripts generated for bioinformatic and statistical analysis are available at https://github.com/magdawutkowska/bistorta.

