## Supplementary materials for Wutkowska et al., Can root-associated fungi mediate the impact of abiotic conditions on the growth of a High Arctic herb? for "Can root-associated fungi mediate the impact of abiotic conditions on the growth of a High Arctic herb?"

**Supplementary 1**

Morphological characteristics of *Bistorta vivipara* measured in this study: rhizome volume (RV; panel A), leaf length (LL; panel B, number 6) and a ratio of inflorescence to stem length (I/S; panel B, ratio of number 2 to 1). Photo: Sunil Mundra.

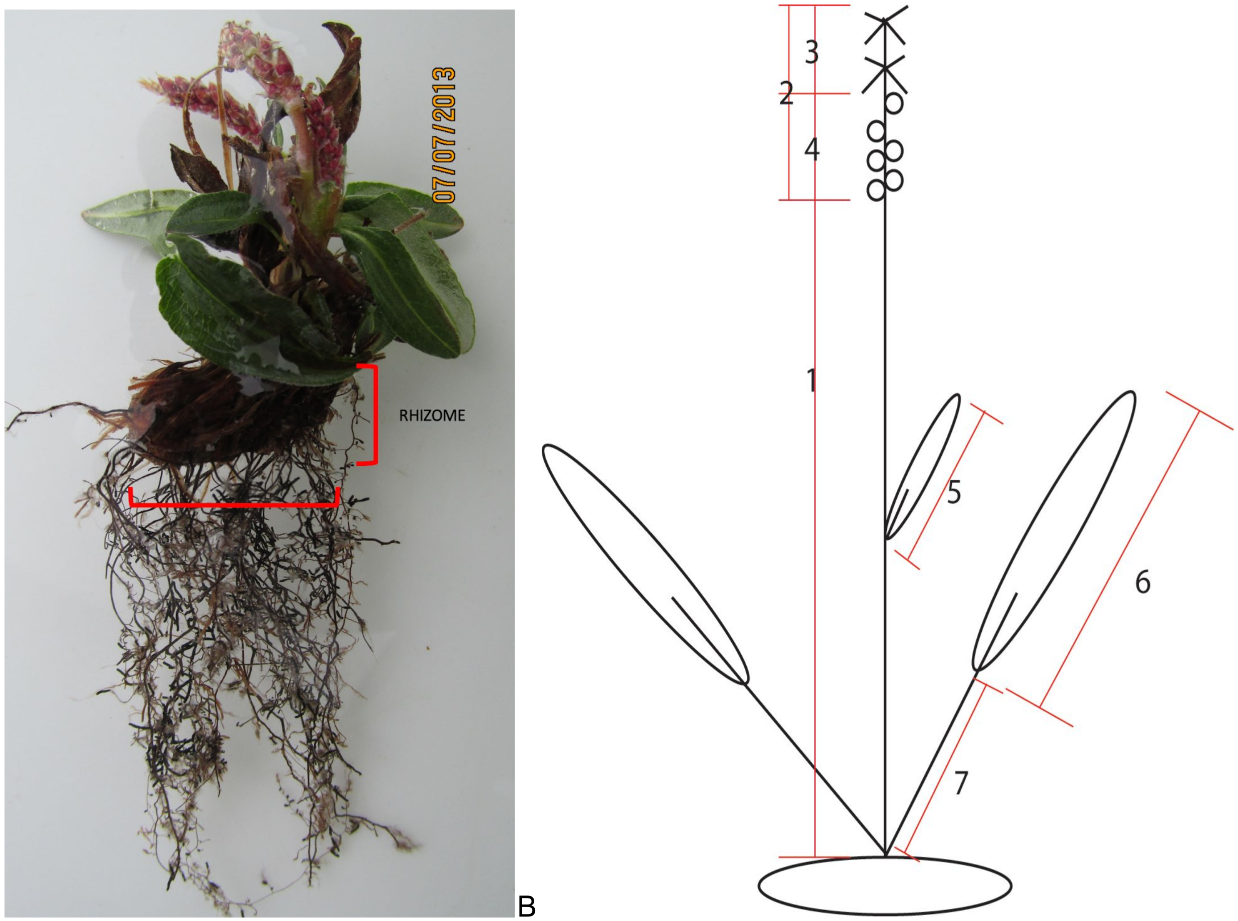


A B

**Supplementary 2**

The overview of bioinformatics pipeline analysing fungal data.

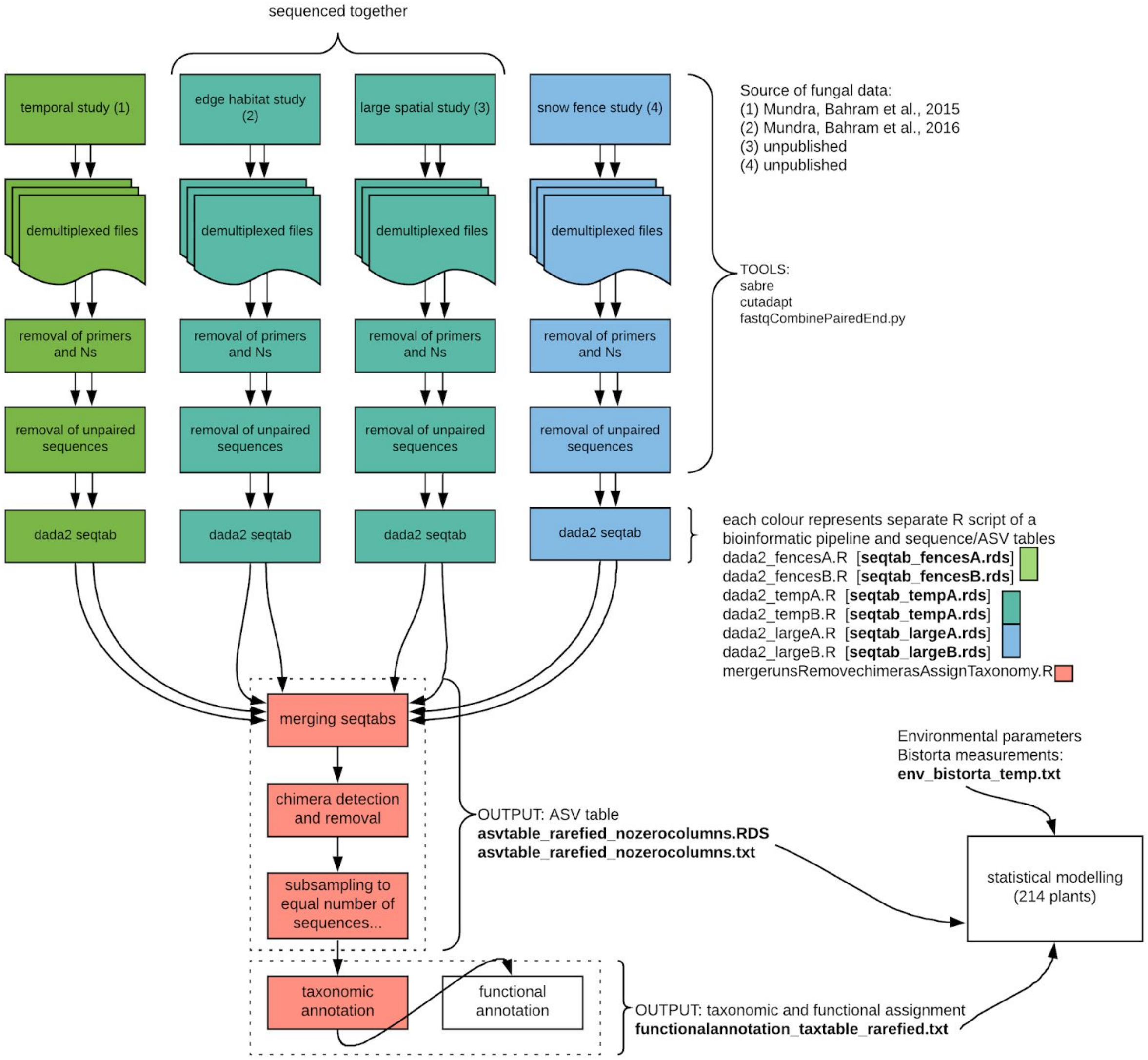

**Supplementary 3**
Characteristics of localities used in this study

| Localities | Description |
|---|---|
| Renardbreen | glacier forefront |
| Hørbyebreen | glacier forefront |
| Trollkjeldene | hot springs |
| Ringhorndalen | arctic steppe |
| Isdammen | natural tundra |
| Vestpynten | nutrient-rich tundra |
| Adventdalen (Snow fences) | natural tundra |
| Bjørndalen (Mine 3 tailings) | nutrient-rich mine-contaminated site |
| Kvalvågen | hydrocarbon-rich site |

a. **How did the localities differ in terms of edaphic variables?**

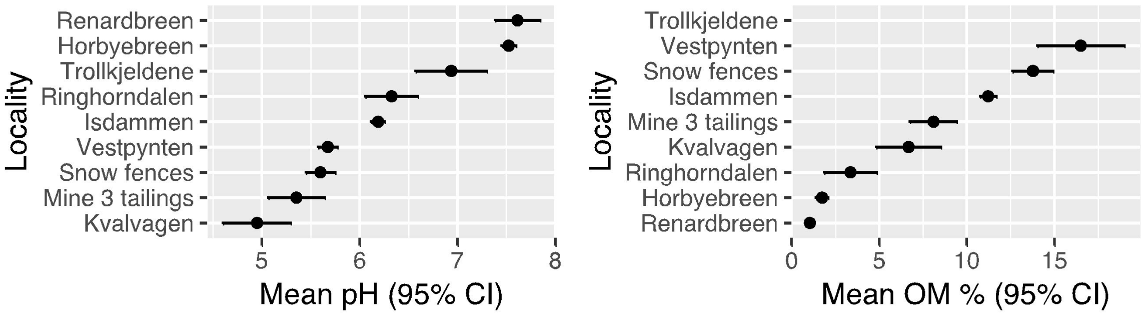

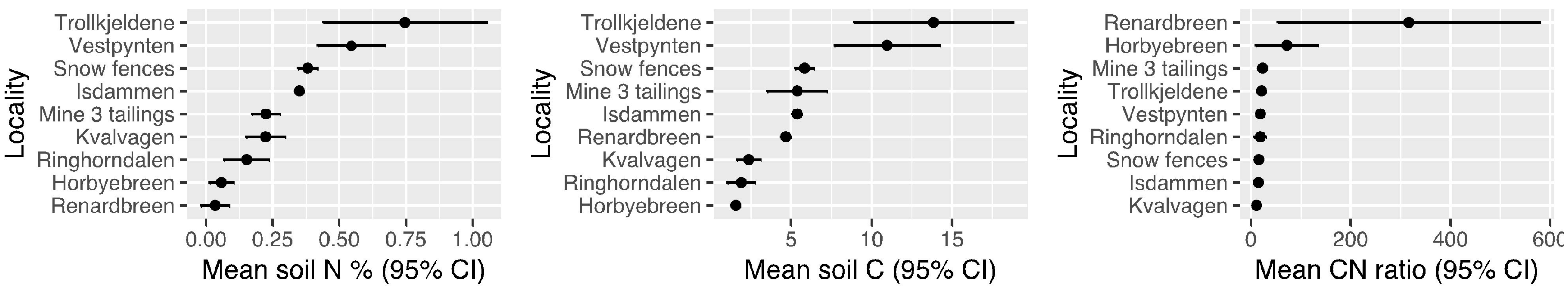


b. **How did plant measurements differ in studied localities?**

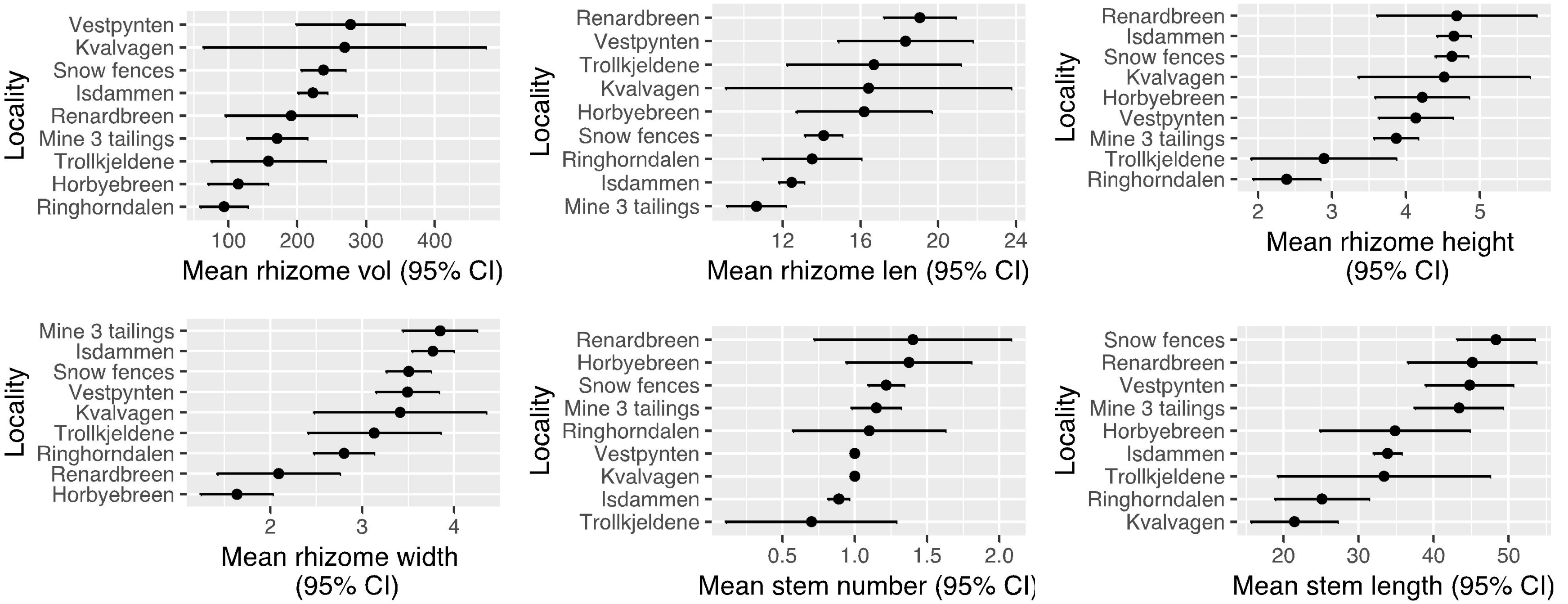

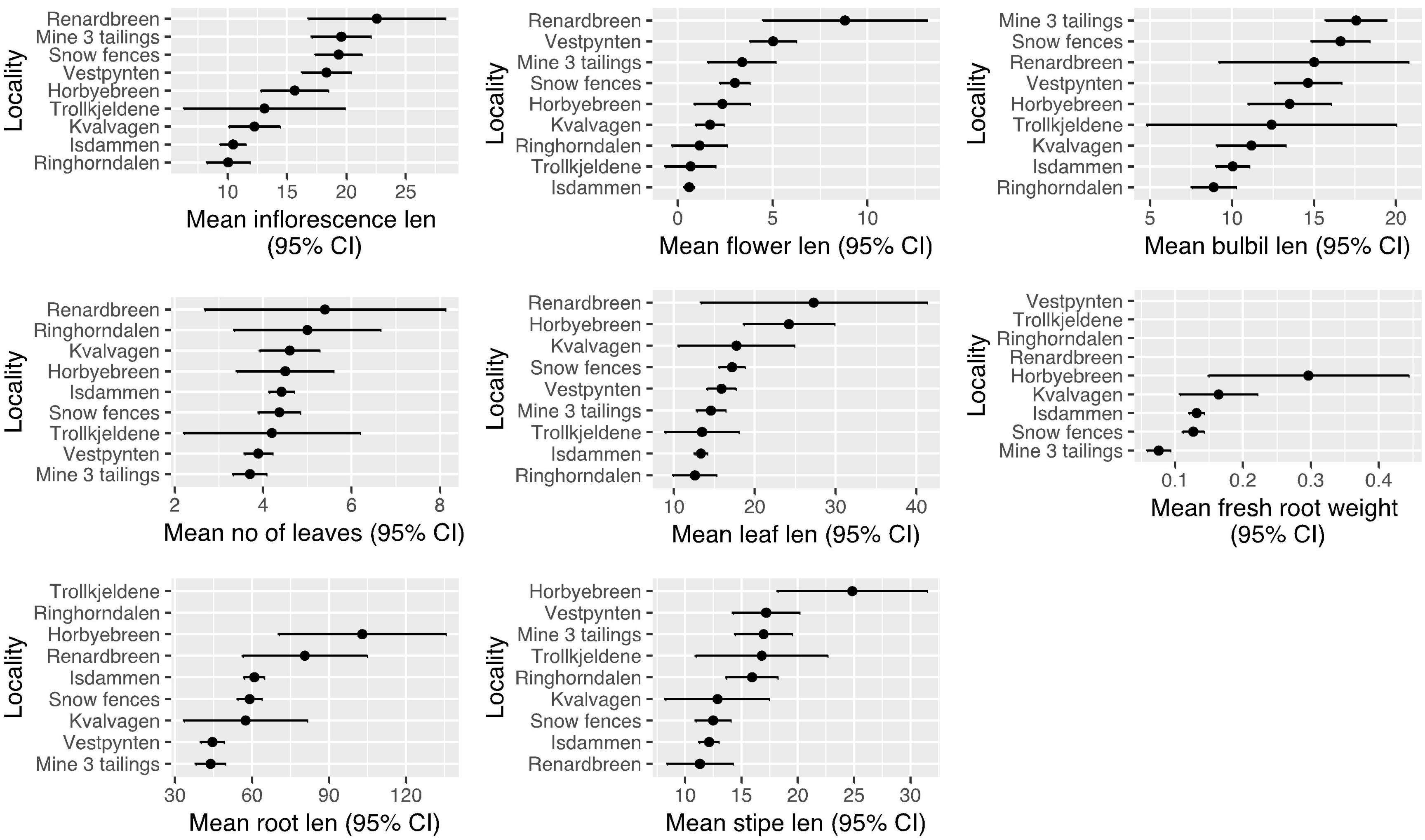
Locality
Renardbreen
Mine 3 tailings
Snow fences
Vestpynten
Horbyebreen
Trollkjeldene
Kvalvagen
Isdammen
Ringhorndalen
10 15 20 25
Mean inflorescence len (95% CI)
Locality
Renardbreen
Vestpynten
Mine 3 tailings
Snow fences
Horbyebreen
Kvalvagen
Ringhorndalen
Trollkjeldene
Isdammen
0 5 10
Mean flower len (95% CI)
Locality
Mine 3 tailings
Snow fences
Renardbreen
Vestpynten
Horbyebreen
Trollkjeldene
Kvalvagen
Isdammen
Ringhorndalen
5 10 15 20
Mean bulbil len (95% CI)
Locality
Renardbreen
Ringhorndalen
Kvalvagen
Horbyebreen
Isdammen
Snow fences
Trollkjeldene
Vestpynten
Mine 3 tailings
2 4 6 8
Mean no of leaves (95% CI)
Locality
Renardbreen
Horbyebreen
Kvalvagen
Snow fences
Vestpynten
Mine 3 tailings
Trollkjeldene
Isdammen
Ringhorndalen
10 20 30 40
Mean leaf len (95% CI)
Locality
Vestpynten
Trollkjeldene
Ringhorndalen
Renardbreen
Horbyebreen
Kvalvagen
Isdammen
Snow fences
Mine 3 tailings
0.1 0.2 0.3 0.4
Mean fresh root weight (95% CI)
Locality
Trollkjeldene
Ringhorndalen
Horbyebreen
Renardbreen
Isdammen
Snow fences
Kvalvagen
Vestpynten
Mine 3 tailings
30 60 90 120
Mean root len (95% CI)
Locality
Horbyebreen
Vestpynten
Mine 3 tailings
Trollkjeldene
Ringhorndalen
Kvalvagen
Snow fences
Isdammen
Renardbreen
10 15 20 25 30
Mean stipe len (95% CI)

c. **How do fungi differ in studied localities?**

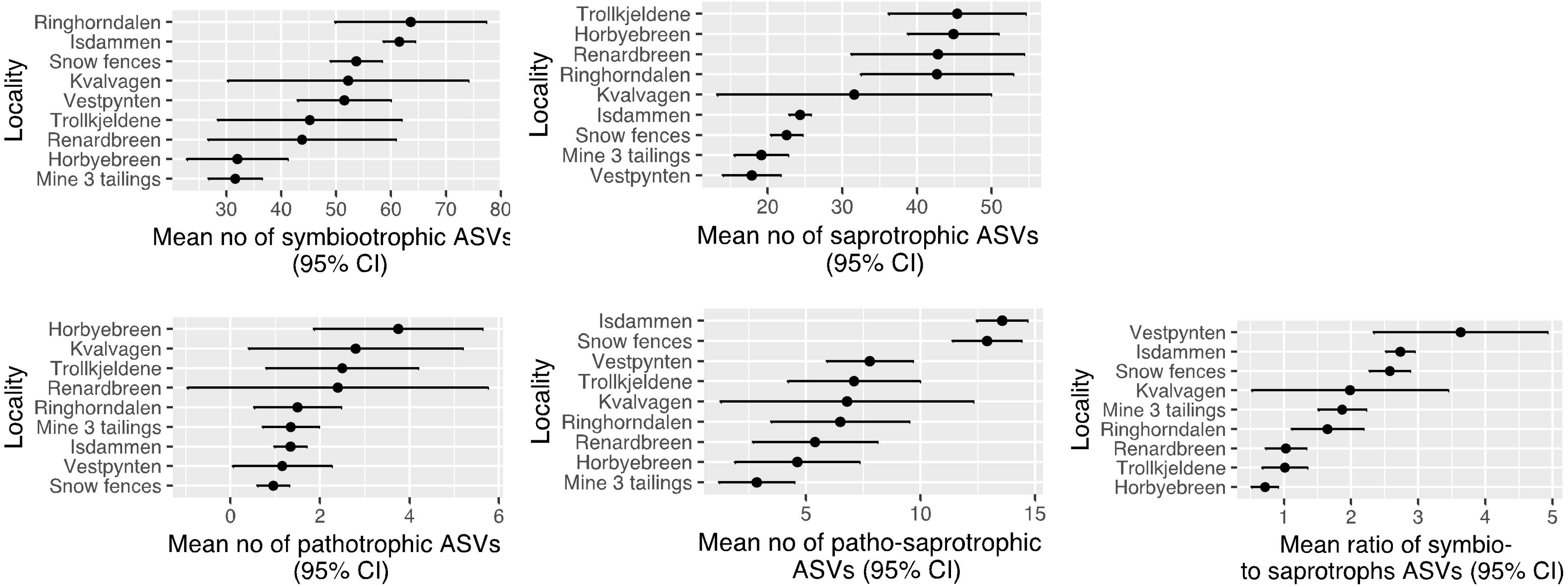

**Supplementary 4**

Complete list of coefficients and estimates calculated in the two best fitting models. Abbreviations: **N** - soil nitrogen content; **CN** - the ratio of soil carbon and nitrogen content; **p** - precipitation; **t** - temperature; **D** - fungal richness (in presence-absence model: number of fungal amplicon sequence variants (ASVs); in abundance model - Shannon-Wiener index); **Sy/Sa** - in presence-absence model: the ratio of symbio- and saprotrophic ASVs and in abundance model: the ratio of symbio- to saprotrophic reads; **CC** - community composition proxy based on presence-absence or read abundance table, respectively; **I/S** - the ratio of inflorescence length to stem length; **RV** - rhizome volume; **LL** - leaf length; statistical significance is coded such as '***' indicate p-value = 0 - 0.001, '**' - 0.001 - 0.01 and '*' 0.01 - 0.05.

a) presence-absence model (Fungal CC not important + no I/S = community composition does not impact plants and no effect of fungi on I/S)

| Response | Predictor | Estimate | Std.Error | DF | Crit.Value | P.Value | Std.Estimate | |
|---|---|---|---|---|---|---|---|---|
| D | N | 0.0980 | 0.1230 | 173 | 0.7967 | 0.4267 | 0.0981 | |
| D | CN | 0.1390 | 0.1151 | 173 | 1.2083 | 0.2286 | 0.1429 | |
| D | pH | -0.0095 | 0.1499 | 173 | -0.0635 | 0.9495 | -0.0066 | |
| D | p | -0.0217 | 0.2350 | 173 | -0.0922 | 0.9266 | -0.0199 | |
| **D** | **t** | **-0.4482** | **0.1266** | **173** | **-3.5410** | **0.0005** | **-0.4048** | *** |
| Sy/Sa | N | 0.0713 | 0.1060 | 173 | 0.6727 | 0.5020 | 0.0723 | |
| Sy/Sa | CN | 0.0376 | 0.0991 | 173 | 0.3789 | 0.7052 | 0.0391 | |
| Sy/Sa | pH | -0.0627 | 0.1292 | 173 | -0.4853 | 0.6281 | -0.0442 | |
| **Sy/Sa** | **p** | **0.4381** | **0.2093** | **173** | **2.0931** | **0.0378** | **0.4072** | * |
| Sy/Sa | t | 0.0382 | 0.1099 | 173 | 0.3476 | 0.7286 | 0.0349 | |
| CC | N | 0.0125 | 0.0679 | 173 | 0.1844 | 0.8539 | 0.0129 | |
| CC | CN | -0.0256 | 0.0636 | 173 | -0.4025 | 0.6878 | -0.0271 | |
| CC | pH | 0.0372 | 0.0829 | 173 | 0.4489 | 0.6541 | 0.0267 | |
| CC | p | 0.4397 | 0.2373 | 173 | 1.8525 | 0.0657 | 0.4147 | |
| **CC** | **t** | **0.2698** | **0.0879** | **173** | **3.0708** | **0.0025** | **0.2504** | ** |
| I/S | N | -0.1020 | 0.1057 | 173 | -0.9644 | 0.3362 | -0.1188 | |
| I/S | CN | -0.0611 | 0.0992 | 173 | -0.6162 | 0.5385 | -0.0731 | |
| I/S | pH | 0.1317 | 0.1292 | 173 | 1.0195 | 0.3094 | 0.1068 | |
| I/S | p | -0.0064 | 0.1726 | 173 | -0.0371 | 0.9704 | -0.0068 | |
| I/S | t | 0.0761 | 0.1062 | 173 | 0.7167 | 0.4746 | 0.0800 | |
| RV | N | 0.0835 | 0.1112 | | 171 | 0.7512 | 0.4535 | 0.0857 |

| | | | | | | | | |
|---|---|---|---|---|---|---|---|---|
| RV | CN | -0.1119 | 0.1046 | 171 | -1.0696 | 0.2863 | -0.1180 | |
| RV | pH | 0.0224 | 0.1362 | 171 | 0.1644 | 0.8696 | 0.0160 | |
| RV | p | 0.2854 | 0.1552 | 171 | 1.8389 | 0.0677 | 0.2686 | |
| **RV** | **t** | **0.2869** | **0.1132** | **171** | **2.5352** | **0.0121** | **0.2656** | * |
| **RV** | **D** | **0.2632** | **0.0716** | **171** | **3.6782** | **0.0003** | **0.2698** | *** |
| RV | Sy/Sa | -0.0983 | 0.0836 | 171 | -1.1756 | 0.2414 | -0.0996 | |
| **LL** | **RV** | **0.5255** | **0.0607** | **170** | **8.6629** | **0.0000** | **0.5252** | *** |
| LL | CN | 0.1178 | 0.0761 | 170 | 1.5477 | 0.1236 | 0.1241 | |
| LL | pH | -0.0283 | 0.0980 | 170 | -0.2889 | 0.7730 | -0.0202 | |
| LL | p | 0.0504 | 0.0896 | 170 | 0.5631 | 0.5741 | 0.0474 | |
| **LL** | **t** | **-0.3429** | **0.0816** | **170** | **-4.2005** | **0.0000** | **-0.3173** | *** |
| LL | D | -0.0063 | 0.0580 | 170 | -0.1086 | 0.9137 | -0.0065 | |
| **LL** | **Sy/Sa** | **-0.1975** | **0.0672** | **170** | **-2.9377** | **0.0038** | **-0.1999** | ** |
| ~~CC | ~~Sy/Sa | 0.1560 | NA | 187 | 2.1428 | 0.0167 | 0.1560 | * |

b) abundance model (no effect of fungi on plants)

| **Response** | **Predictor** | **Estimate** | **Std.Error** | **DF** | **Crit.Value** | **P.Value** | **Std.Estimate** | |
|---|---|---|---|---|---|---|---|---|
| **D** | **N** | **0.2355** | **0.1108** | **171** | **2.1258** | **0.0350** | **0.2470** | * |
| D | CN | -0.0124 | 0.1043 | 171 | -0.1190 | 0.9054 | -0.0134 | |
| D | pH | -0.1292 | 0.1366 | 171 | -0.9459 | 0.3455 | -0.0944 | |
| D | p | -0.0515 | 0.1670 | 171 | -0.3083 | 0.7582 | -0.0496 | |
| D | t | -0.0288 | 0.1105 | 171 | -0.2607 | 0.7946 | -0.0273 | |
| **Sy/Sa** | **N** | **-0.2792** | **0.1015** | **171** | **-2.7513** | **0.0066** | **-0.2967** | ** |
| **Sy/Sa** | **CN** | **-0.2012** | **0.0973** | **171** | **-2.0686** | **0.0401** | **-0.2199** | * |
| Sy/Sa | pH | 0.2166 | 0.1239 | 171 | 1.7482 | 0.0822 | 0.1603 | |
| Sy/Sa | p | 0.1372 | 0.1084 | 171 | 1.2654 | 0.2075 | 0.1338 | |
| Sy/Sa | t | 0.0542 | 0.1014 | 171 | 0.5349 | 0.5934 | 0.0521 | |
| CC | N | -0.0341 | 0.0844 | 171 | -0.4039 | 0.6868 | -0.0347 | |

| | | | | | | | | |
|---|---|---|---|---|---|---|---|---|
| CC | CN | -0.0165 | 0.0790 | 171 | -0.2087 | 0.8349 | -0.0173 | |
| CC | pH | -0.2032 | 0.1037 | 171 | -1.9595 | 0.0517 | -0.1442 | |
| CC | p | 0.1709 | 0.1979 | 171 | 0.8634 | 0.3891 | 0.1598 | |
| **CC** | **t** | **0.3168** | **0.0917** | **171** | **3.4558** | **0.0007** | **0.2916** | *** |
| I/S | N | -0.1016 | 0.1059 | 171 | -0.9598 | 0.3385 | -0.1184 | |
| I/S | CN | -0.0572 | 0.0995 | 171 | -0.5754 | 0.5658 | -0.0686 | |
| I/S | pH | 0.1153 | 0.1304 | 171 | 0.8846 | 0.3776 | 0.0936 | |
| I/S | p | -0.0119 | 0.1716 | 171 | -0.0694 | 0.9448 | -0.0127 | |
| I/S | t | 0.0734 | 0.1062 | 171 | 0.6913 | 0.4903 | 0.0773 | |
| RV | N | 0.1342 | 0.1084 | 171 | 1.2383 | 0.2173 | 0.1385 | |
| RV | CN | -0.0520 | 0.1034 | 171 | -0.5034 | 0.6153 | -0.0553 | |
| RV | pH | 0.0183 | 0.1344 | 171 | 0.1359 | 0.8921 | 0.0131 | |
| RV | p | 0.2171 | 0.1324 | 171 | 1.6395 | 0.1029 | 0.2056 | |
| RV | t | 0.1636 | 0.1081 | 171 | 1.5131 | 0.1321 | 0.1526 | |
| **LL** | **RV** | **0.5431** | **0.0603** | **170** | **9.0090** | **0.0000** | **0.5395** | *** |
| **LL** | **N** | **-0.2272** | **0.0799** | **170** | **-2.8434** | **0.0050** | **-0.2331** | ** |
| LL | CN | 0.1374 | 0.0756 | 170 | 1.8167 | 0.0710 | 0.1449 | |
| LL | pH | 0.0113 | 0.0953 | 170 | 0.1184 | 0.9059 | 0.0081 | |
| LL | p | -0.0245 | 0.0822 | 170 | -0.2978 | 0.7662 | -0.0230 | |
| **LL** | **t** | **-0.3739** | **0.0788** | **170** | **-4.7443** | **0.0000** | **-0.3464** | *** |
| ~~Sy/Sa | ~~D | -0.4151 | NA | 185 | -6.1557 | 0.0000 | -0.4151 | *** |
